# Dissection of molecular assembly dynamics by tracking orientation and position of single molecules in live cells

**DOI:** 10.1101/068767

**Authors:** Shalin B. Mehta, Molly McQuilken, Patrick La Riviere, Patricia Occhipinti, Amitabh Verma, Rudolf Oldenbourg, Amy S. Gladfelter, Tomomi Tani

**Affiliations:** Eugene Bell Center for Regenerative Biology and Tissue Engineering, Marine Biological Laboratory, Woods Hole, MA02543; Department of Biological Sciences, Dartmouth College, Hanover, NH03755; Department of Radiology, University of Chicago, Chicago, IL60637; Physics Department, Brown University, Providence, RI02912; Department of Biology, University of North Carolina at Chapel Hill, Chapel Hill, NC 27599

**Keywords:** single molecule orientation, live cell imaging, polarized fluorescence, actin, septin

## Abstract

Regulation of order, such as orientation and conformation, drives the function of most molecular assemblies in living cells, yet remains difficult to measure accurately through space and time. We built an instantaneous fluorescence polarization microscope, which simultaneously images position and orientation of fluorophores in living cells with single-molecule sensitivity and a time resolution of 100ms. We developed image acquisition and analysis methods to track single particles that interact with higher-order assemblies of molecules. We tracked the fluctuations in position and orientation of molecules from the level of an ensemble of fluorophores down to single fluorophores. We tested our system *in vitro* using fluorescently labeled DNA and F-actin in which the ensemble orientation of polarized fluorescence is known. We then tracked the orientation of sparsely labeled F-actin network at the leading edge of migrating human keratinocytes, revealing the anisotropic distribution of actin filaments relative to the local retrograde flow of the F-actin network. Additionally, we analyzed the position and orientation of septin-GFP molecules incorporated in septin bundles in growing hyphae of a filamentous fungus. Our data indicate that septin-GFP molecules undergo positional fluctuations within, ∼350nm of the binding site and angular fluctuations within ∼30^°^ of the central orientation of the bundle. By reporting position and orientation of molecules while they form dynamic higher-order structures, our approach can provide new insights into how micron-scale ordered assemblies emerge from nanoscale molecules in living cells.

**Significance Statement:** In living cells, the 3D architecture of molecular assemblies such as chromosomes, lipid bilayers, and the cytoskeleton is regulated through the interaction among their component molecules. Monitoring the position and orientation of constituent molecules is important for understanding the mechanisms that govern the structure and function of these assemblies. We have developed an instantaneous fluorescence polarization microscope to track the position and orientation of fluorescently labeled particles, including single molecules, which form micron-scale macromolecular assemblies in living cells. Our imaging approach is broadly applicable to the study of dynamic molecular interactions that underpin the function of micron-scale assemblies in living cells.

## Introduction

The generation and dissolution of order within populations of biological molecules is central to almost all cellular processes. The emergence of ordered arrays of biological molecules is manifested in lipid membranes, DNA, the cytoskeleton, and many other molecular assemblies in the cytoplasm of the living cell. Polarization-resolved fluorescence imaging of densely labeled assemblies has successfully probed their net order and architectural dynamics (1–5). In addition, tracking of sparsely-labeled cytoskeletal networks (6–9) has illuminated how turnover of individual molecules enables transitions in the spatial organization of cytoskeletal networks. We have combined tracking of both orientation and position of single fluorophores associated with cystoskeletal proteins in live cells to analyze the dynamics of cytoskeleton assembly.

Fluorophores, including the green fluorescent protein (GFP), emit fluorescence by radiating light as dipoles. The light emitted from a single dipole is fully polarized, with most of the energy polarized along the dipole axis. For example, the absorption and emission dipole axes of GFP chromophores are fixed within the GFP molecule, as was found in GFP crystals (10, 11). Therefore, fluorescent labels can report the orientation of biomolecules as long as the labels are rigidly bound to the biomolecules, constraining their relative motion (12, 13). Comparison of the intensities recorded either by polarization-resolved excitation (3–5, 14–18) or polarization-resolved detection (1, 2, 19–21) has allowed measurement of orientations of fluorophores.

For fluorescent assemblies that exhibit simple geometries, such as planar (3), spherical (2), or cylindrical (19) shapes, two orthogonal polarization-resolved measurements suffice to retrieve fluorophore orientations relative to these geometries. However, unbiased measurement of dipole orientation in a complex assembly requires exciting or analyzing the fluorescent dipoles using at least three and preferably four polarizer orientations (22). We have previously developed (4, 5, 14, 15) a liquid crystal (LC) based fluorescence polarization microscope (fluorescence LC-PolScope) that sequentially excites fluorophores with light polarized along 0^,°^, 45^°^, 90^°^ and 135^°^. We used the fluorescence LC-PolScope to analyze the remodeling of septin assemblies with a time resolution of about 10 seconds (4, 5). For assemblies that rearrange on a faster time scale, the fluorescence LC-PolScope generates spurious results because of motion artifacts introduced by sequential image acquisition. Simultaneous acquisition of polarization-resolved images circumvents the artifacts introduced by sequential acquisition.

Simultaneous analysis of fluorescence emission along two orthogonal orientations was recently combined with localization microscopy to reveal the angular distribution of single fluorescent dipoles attached to filamentous assemblies in fixed cells and *in vitro* (19). For live cell imaging, however, localization microscopy is typically too slow. Further, interpretation of angular distribution from two orthogonally polarized images requires a subtle, yet important, assumption that all fluorophores in the analyzed puncta are bound to the same filament of cylindrical symmetry. This assumption often does not hold in live cells either due to the complexity of the ordered structure (e.g., F-actin network at lamellipodia or septin hourglass at yeast bud-neck) or non-cylindrical symmetry of the structure (e.g. focal adhesions). Widefield unpolarized excitation and simultaneous analysis of fluorescence emission along four polarization orientations (20) has been used to study 3D rotation of isolated myosin motor proteins walking on actin filaments, but not applied to micron-scale assemblies in living cells that display interactions between many molecules. Thus, previous approaches are either incompatible with live cell imaging or limited in their capacity to track position and orientation of many molecules in parallel against the fluorescent background typical of a live cell.

We have developed an instantaneous fluorescence polarizing microscope (instantaneous FluoPolScope) that combines isotropic total internal reflection (TIR) excitation and instantaneous sorting of emitted fluorescence along four polarization orientations using a home-built image-splitting device. These optics are complemented by algorithms for tracking of position and orientation of fluorescent particles, including single molecules, at the cortex of live cells with sub-pixel accuracy. Our method has the potential to answer important questions raised by studies using high-resolution X-ray crystallography, NMR or electron microscopy about dynamic molecular interactions during assembly/disassembly of biological molecules in living cells or in systems reconstituted *in vitro*.

We report two approaches of optically dissecting the ordered organization of micron-scale molecular assemblies in live cells: 1) Tracking the position and orientation of molecular components within complex networks that are sparsely labeled with fluorescent orientation probes. This approach can reveal the dynamic rearrangement of the molecular components associated with the deformation or movement of the network. 2) Tracking the position and orientation of fluorescently labeled protein subunits that interact with a higher-order assembly, thus revealing their molecular interactions and assembly kinetics.

For establishing and testing the new methods, we observed fluorescently labeled DNA and actin filaments prepared *in vitro* and bound to a coverslip surface. We evaluated the utility of the methods to study molecular order of micron-scale assemblies of actin and septin in live cells. We used sparse labeling of F-actin at the leading edge of migrating human keratinocytes to analyze changes in the orientation of actin filaments relative to the direction of actin retrograde flow. We expressed septins with rotationally constrained GFP (conGFP) tags (4, 5, 12, 13) in the filamentous fungus *Ashbya gossypii*, and observed the change in polarized fluorescence of individual septin particles and their assemblies located at the cortex of living cells. Tracking temporal fluctuations of the fluorescence intensity, position, and polarization of septin particles revealed confined motion, both in position and in angle, experienced by septin-GFP molecules, thus providing new insight in their assembly kinetics in living cells.

## Results and Discussion

### Instantaneous imaging of position and orientation of fluorophores with single-molecule sensitivity

We designed a microscope to provide sub-pixel position and bias-free measurement of molecular orientation of single dipoles and their ensembles near a glass-water interface or at the cell cortex. The angular distribution of dipoles (Fig. 1A) is represented by its azimuth *ϕ* within the focal plane (XY), tilt *θ* relative to the optical axis (Z), and a wobble range *δ* over the camera exposure. Polarization analysis in the image plane provides unambiguous measurement of the azimuth, called orientation in this paper. However, the tilt and the wobble are both inferred from the polarization factor *p*. The polarization factor varies from 0 (isotropic) to 1 (fully anisotropic) depending on the tilt and wobble in the distribution of fluorophores.

**Fig. 1.**
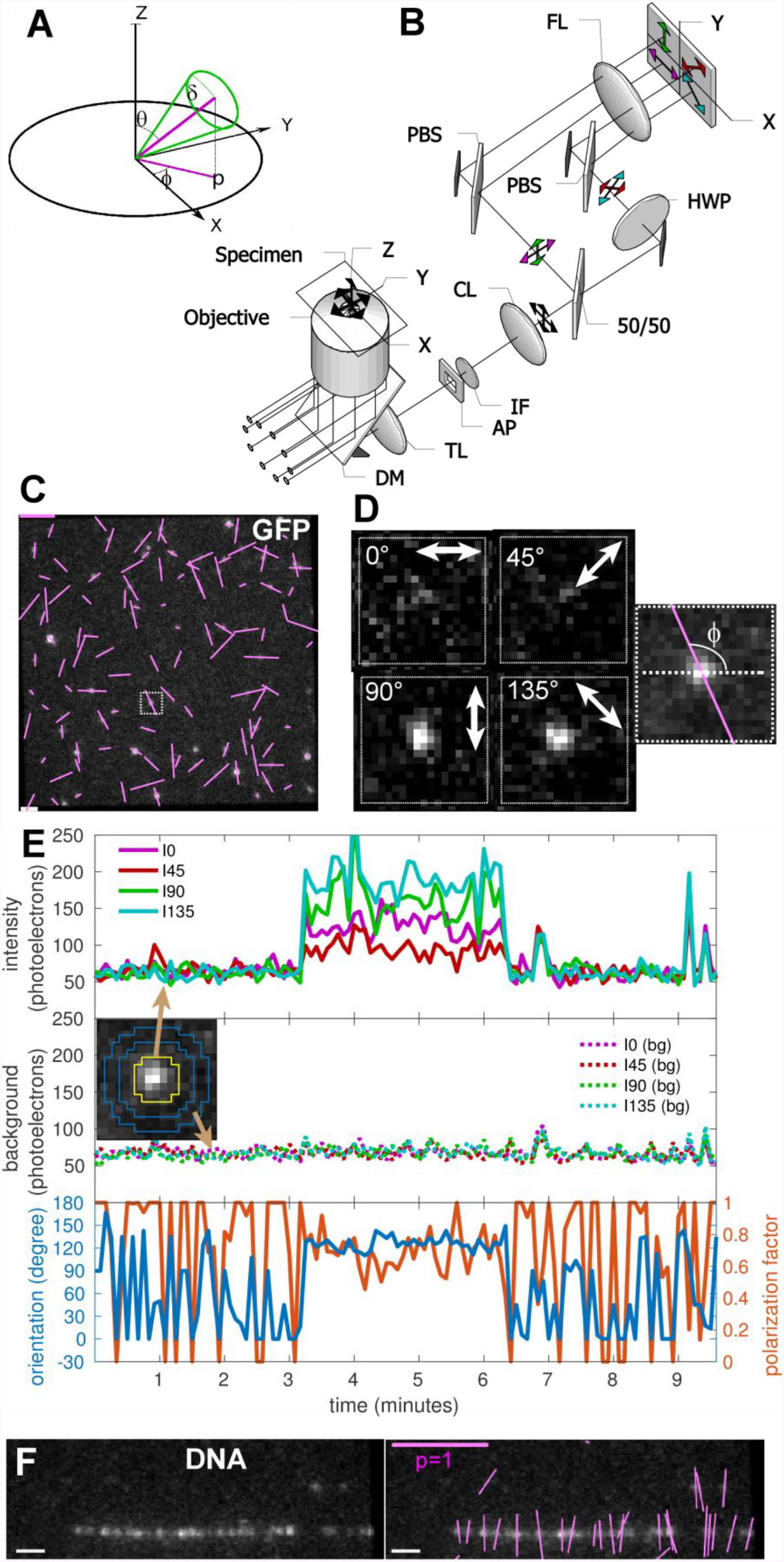
Imaging orientation of fluorescent single molecules with instantaneous FluoPolScope: (**A**) Coordinate system used to parameterize single dipoles and their ensembles (thick green line through origin) by their net orientation *ϕ* in the focal plane (XY), net tilt *θ* relative to the optical axis of the microscope (Z), and wobble *δ* during the exposure. (**B**) Schematic of the microscope. (**C**) Polarization resolved fluorescence images of GFP molecules attached to coverslip. **(D)** Enlarged images of single GFP molecule identified by outlined square in C. The right-most image shows the computed dipole orientation as magenta line overlaid on the sum of all polarization-resolved images. (**E**) Particle intensities (top), background intensities (middle), and orientation and polarization factor (bottom) of the single GFP molecule shown in D that was recorded for more than 3 minutes. The inset in the middle graph identifies the pixels used to estimate particle intensities (area enclosed by yellow line) and background intensity (circular area between two blue bands) in each quadrant. (**F**) Fluorescence images of λ phage DNA stained with 50 nM of TOTO-1 (left) and the image with orientations of polarized fluorescence of detected particles shown by magenta lines (right). Scale bars: white (F) is 1μm, magenta (C) is polarization factor=1.

For efficient excitation of dipoles oriented in 3D, we used isotropically-polarized total internal reflection (TIR) illumination (Fig. 1B, Methods). For instantaneous imaging of azimuth and polarization factor, we created an imaging path with beam splitters projecting four polarization-resolved images of the field of view onto the four quadrants of an EMCCD camera (Fig. 1B, Methods). We identified the linear, shot-noise limited operating regime of our EMCCD camera using photon transfer analysis (SI Note 1, Fig. S1), which also allows us to express all intensity measurements in the units of detected photoelectrons. Before computing orientation and polarization factors from quadrant intensities, differences in spatial registration and transmission efficiencies of the quadrants are accounted for using a robust calibration system (Methods, Fig. S2, and SI Note 2). In utilizing the calibration, we expressed the orientation of polarized fluorescence, polarization factor, and intensity in terms of the Stokes parameters of the fluorescence emission in the image plane (Methods, Eqs. 1-3).

To evaluate the sensitivity to single molecules, we imaged monomeric EGFP molecules (Fig. 1C) attached to poly-D-lysine coated glass coverslips (SI Note 3). Unlike the isotropic fluorescence of 100nm beads labeled with a large number of fluorophores (Fig. S2A), we could detect anisotropic fluorescence of single EGFP molecules by a clear imbalance of their florescence intensities in the four quadrants (Fig. 1D). Imaging with low laser power of 1.1 μW/μm^2^ at the specimen plane (20% ND filter) and 500ms exposure at 5s intervals, we were able to image orientations of single immobile EGFP molecules for up to 15 minutes and longer.

We developed MATLAB code for automated detection of particle locations, intensities, and background (SI Note 4, Fig. S4) to map temporal fluctuations in intensity and orientation of fluorescent particles (Fig. 1E) across the field of view (Figs. 1C, S3A). The fluorescence recorded in the four quadrants from the particle illustrated in Figs. 1D and 1E maintained their relative intensities for up to 3 minutes before simultaneously bleaching to the background level in a single step. EGFP particles were classified as being single molecules bound to the glass or to cellular structures later in the paper by two criteria: 1) their fluorescence resided within a diffraction-limited spot that could be tracked for at least 5 consecutive frames, and 2) their fluorescence bleached to the background intensity in a single step (Video S1).

The nearly uniform histogram of orientations of more than 7000 single EGFP particles (Fig. S3B) confirms that our imaging and calibration approach indeed provides bias-free measurement of orientation. Sum of quadrant intensities above the background (Fig. S3C) exhibits a single peak at ~350 photoelectrons for single EGFP molecules with lognormal distribution typically observed with TIRF illumination (23). Plotting the fluctuations in the detected centroids of the EGFP particles (Fig. S3D) showed that many of them were confined to an area of 25nmx25nm, which is an upper bound of the achieved localization precision.

To estimate the shot-noise induced standard deviation σ_ϕ_ of measured orientations, we propagated the shot-noise of the intensities and expressed it in terms of the sum intensity above background, background intensity, and polarization factor (SI Notes 5 and 6). Their functional dependence is shown in Fig. S5. For the EGFP molecule shown in Figs. 1D and 1E [*p*≈0.8, *I*(total)≈275, *Ibg*(total)≈200], the standard deviation of estimated orientation is 3^°^, which is better than the 10^°^ accuracy reported earlier with a similar polarization analysis system (20) or with a system that analyzes the shape of the emission pattern at a slightly out-of-focused plane (24). The 3^°^ standard deviation is comparable to the accuracy achieved when analyzing the emission pattern at the back focal plane (25).

We used combed λ-phage DNA labeled with TOTO-1 (Fig. 1F, SI Note 3) in which the dipole orientation of the chromophores in the complex has already been characterized (26). As expected, the fluorescence polarization of a TOTO-1 particle was found to be perpendicular to the local orientation of the DNA (Fig.1F, Video S2). We tracked (27) the time-dependent position, intensity, orientation, and polarization factor of sub-resolution particles (SI Note 7, Figs. S6C-G) with the following results: During the observation, the intensity of individual TOTO-1 particles decreased in a step-wise manner, before bleaching down to the background level (Fig. S6E). The lowest intensity step was ~150 photoelectrons. The fluorescence intensity histogram of TOTO-1 particles that bleach to background during acquisition (SI Note 8) showed a broad distribution ranging from 50 to 1500 photoelectrons (Fig. S6H) with periodic peaks at multiples of 150 photoelectrons, which corroborates the single chromophore intensity before complete bleaching (Fig. S6E). The orientation of this TOTO-1 particle (Fig. S6F) shows a stable position and orientation perpendicular to the DNA axis. However, at the single chromophore level, the orientation, position, and polarization factor of the particle (Figs. S6C-G) became more variable, presumably because of a break in the DNA due to a photochemical reaction where TOTO-1 is bound.

Above results show that our microscope can provide a robust and instantaneous measurement of intensity, orientation, and position of single dipoles and their ensembles incorporated in micron-scale assemblies.

### Tracking position and orientation of actin assembly *in vitro* and in living cells

Migratory cells rely on the regulated polymerization of actin networks and the contractile behavior of actomyosin assemblies to generate propulsive forces. The molecular architecture of the actin network at the leading edge and its spatio-temporal evolution has been studied extensively using electron microscopy (28, 29) and fluorescent speckle microscopy (7–9). We used our instantaneous FluoPolScope and tracking algorithms to analyze local orientations of constituent actin filaments relative to retrograde flow of the actin network.

We first identified the orientation of polarized fluorescence of Alexa Fluor 488 phalloidin (AF488-phalloidin, Molecular Probes) bound to *in vitro* actin filaments at different labeling ratios ranging from dense labeling to single molecules of AF488. When exposed to low concentrations (2nM - 5nM, SI Note 3) of AF488-phalloidin, *in vitro* actin filaments were sparsely labeled (Fig. 2A), leading to sub-resolution particles, whose position, fluorescence intensity, orientation, and polarization factor were tracked over time (SI Notes 7 and 8), with the following results.

**Fig. 2.**
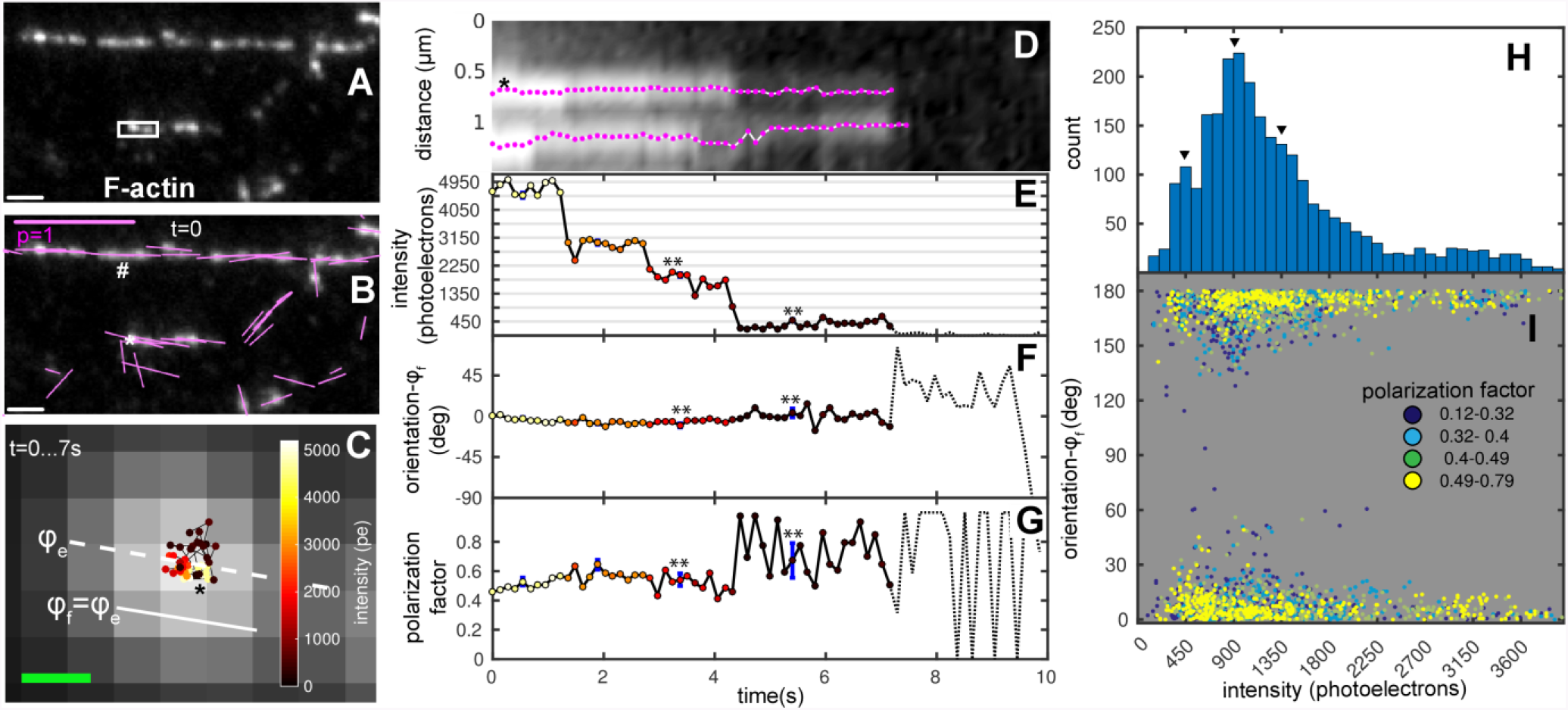
Position and orientation of Alexa Fluor 488 phalloidin bound to F-actin *in vitro*: (**A**) Fluorescence image of F-actin stained with Alexa Fluor (AF) 488 phalloidin (2nM in actin polymerization buffer) at the start of image sequence. **(B)** The image as in (A) with orientations of polarized fluorescence shown by magenta lines. (#)Measured orientation aligns with a straight stretch of the filament. (*)Representative particle used to illustrate tracking of position, intensity, orientation, and polarization factor. (**C**) Positions of centroid of representative particle (* in B) over time connected by a black line. Hue of the dots represents the particle intensity (see color map in inset). Centroids are overlaid on a magnified, averaged fluorescence image from the start to the end of the track. Dashed white line shows ensemble orientation (*ϕ_e_*) of polarized fluorescence of the pixels comprising the particle over the first 3 frames. The ensemble orientation is used to define the local filament orientation (*ϕ_f_*). (**D**) Kymograph of fluorescence particles along the long axis of the white rectangle in A. Position of the particles in the filament is plotted in the vertical direction and time along the horizontal direction. The tracks of 2 particles are shown in magenta. * is the representative particle identified in B,C. (**E**) Time dependent intensity change of the representative particle. (**F**) Time dependent orientation change of the representative particle relative to local filament orientation *ϕ_f_*. ** When the ensemble reduces to a single fluorophore of AF488, its orientation remains close to the filament orientation. (**G**) Polarization factor of the representative particle. ** The polarization factor increases and becomes noisier when the ensemble reduces to single fluorophore. In E, F and G, the dotted line shows the intensity, orientation and polarization factor after the track bleaches to the background, respectively. The uncertainty (± s.d.) in the measured quantities due to shot noise is shown by blue error bars at chosen time points. In some graphs, blue error bars are too small to be clearly visible. (**H**) Histogram of particle intensities over ~2600 individual observations (60 tracked particles). Peaks at multiples of 450 photoelectrons (arrowheads) suggest the intensity of a single AF488. (**I**) Scatter plot of polarization orientation relative to filament axis vs. intensity for the same observations. Individual scatter points are colored by their polarization factor *p* according to legend. Total number of scatter points corresponds to the same number of individual observations in H. Scale bars: white (B) is 1μm, magenta (B) is polarization factor=1, green (C) is 100nm.

The ensemble orientation of fully decorated straight filaments was found to be parallel to the orientation of the filament axis (# in Fig. 2B). Actin filaments exhibit bends, making it difficult to identify filament orientation at the location of each sub-resolution particle. Therefore, we defined the local filament orientation (ϕ*_f_*) (Fig. 2C) as the ensemble orientation (ϕ*_e_*) averaged over the first five time-points of tracked particles. We observed displacements of tracked particles within a range of ~100nm (Figs. 2C, 2D, Video S3), which is well above our experimentally verified localization accuracy of 25nm (Fig. S3D). This displacement of AF488-phalloidin particles can be attributed to the flexing of actin filaments (30). The fluorescence intensity of AF488-phalloidin particles typically exhibited several bleaching steps before their intensity was reduced to the background level (Fig. 2E). The minimum step size was ~450 photoelectrons. The fluorescence intensity histogram of particles tracked from ensemble to no fluorophores (Fig. 2H) shows peaks at multiples of ~450 photoelectrons, which corroborate the intensity of single AF488 as shown in Fig. 2E. The orientation of AF488-phalloidin was consistently parallel to the filaments throughout our observations (Fig. 2F), even at the single molecule intensity level. Interestingly, for several AF488 particles, we observed that the polarization factor remained high or even increased (Fig. 2G) when the fluorophore ensemble bleached to a single fluorophore. These observations are confirmed by plotting the particle orientation (relative to local filament orientation) against the particle intensity with color-coded polarization factor in Fig. 2I, where the majority of particles with single molecule intensity (~450 photoelectrons) showed polarization factors above 0.49 and the particles with high polarization factor reported orientation of the filament irrespective of the intensity. Data in Figs. 2F and 2I strongly suggests that the fluorescence polarization of AF488-phalloidin faithfully reports the orientation of actin filaments, even at the single molecule level.

Our method, therefore, offers the possibility of simultaneously analyzing the structural dynamics of actin networks in live cells and the molecular orientation of the constituent actin filaments that are sparsely labeled with AF488-phalloidin. We introduced low concentrations (10-20 nM) of AF488-phalloidin to human keratinocytes (HaCaT cells) cultured on glass coverslips (SI Note 9). Sparse labeling of F-actin in live cells with fluorescent phalloidin at concentrations lower than ~20 nM preserves normal actin driven dynamics and cell motility in variety of cell types (8, 31), enabling visualization of continuous retrograde flow within actin networks at the leading edge.

We observed distinct particles of AF488-phalloidin across the leading edge (Video S4 and Video S5) of migrating HaCaT cells. For further analysis, we selected cells that showed periodical protrusion and retraction of lamellipodia at the leading edge even after introducing fluorescent phalloidin into the cytoplasm (Fig. 3, Video S4). Sparse fluorescent labeling allowed us to track the position, motion, and orientation of individual particles. Using maximum intensity projections of the particle fluorescence over time, we confirmed that particle tracks fully probed the actin network at the leading edge and its movement to the base of the lamellipodium (Fig. 3A). We tracked the flow direction of each AF488-phalloidin particle (Figs. 3B) and compared it to the orientation of the associated actin filaments measured by its fluorescence polarization. We monitored the changes in the local orientation of F-actin (Fig. 3C) during the continuous retrograde flow (Fig. 3D) within two domains of the actin network: the distal domain at the leading edge and the proximal domain of the lamella consisting of actin arcs (7, 9).

**Fig. 3.**
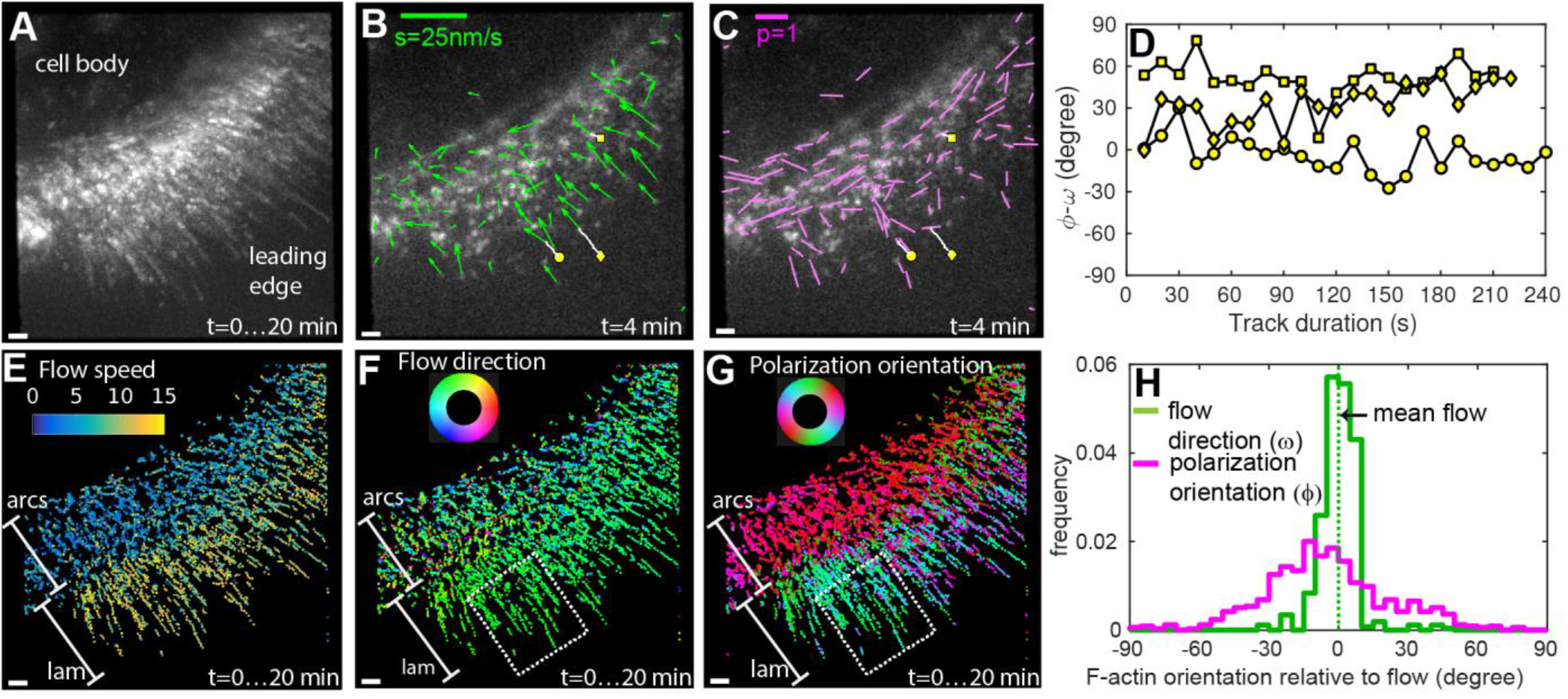
Position and orientation of Alexa Fluor 488 phalloidin bound to F-actin during retrograde flow in live cells: **(A)**Maximum intensity projection of F-actin network sparsely labeled with AF488 phalloidin at the leading edge of a migrating human keratinocyte (HaCaT cell). Maximum intensity projection over the 20 minutes long movie shows complete coverage of the leading edge consisting of actin arcs and lamellipodium. **(B)** The direction (ω) and the speed (*s*) of the movement estimated from the motion of the tracked particle are shown by green arrows. The direction of the arrow indicates the local direction of the movement and the length of the arrow indicates the speed. Trajectories of a few fluorescent particles are shown by white lines. **(C)** Orientation of polarized fluorescence (ϕ) and polarization factor (*p*) are shown by magenta lines. Orientation of the lines represents the polarization orientation, and the length of the lines represents the polarization factors. **(B & C)** Yellow circle, square, and diamond mark the starting positions of three tracks plotted in (D), whereas the white lines indicate the trajectory of the particle up to the chosen time point of t=4min. **(D)** Time-resolved fluctuations in the fluorescence polarization orientation of individual particles tracked in (B & C). The orientation is plotted relative to the net direction of the track computed by vector averaging of all flow vectors along the track. **(E)** Temporal projection of flow speed is shown as a scatter plot of dots placed at their detected positions and colored according to their local retrograde flow speed. The color is assigned according to color scale on top-left. **(F)** Temporal projection of moving directions is constructed as described in (E), but using the color-wheel with periodicity of 360 shown on top-left. **(G)** Temporal projection of orientation of polarized fluorescence constructed as described in (E), but using the color-wheel with periodicity of 180 shown on top-left. **(F & G)** The dashed white line identifies the region of interest used for statistical analysis in (H). **(E,F, &G)** Bands parallel to leading edge corresponding to actin arcs are marked as *arcs* and the band corresponding to lamellipodium is marked as *lam*. **(H)** Normalized histogram of retrograde flow directions (green line) and polarization orientations (magenta line) of AF488 phalloidin particles in the area enclosed by white dashed line in (F) and (G). The distribution of retrograde flow direction and polarization orientation is plotted relative to the mean retrograde flow direction (dashed green line) within the region of interest. Scale bars: white (A-C,E-G) is 1μm, magenta (C) is polarization factor=1, green (B) is flow speed (*s*)=25nm/s.

Spatial maps of flow speed (Fig. 3E, Fig. S7A), flow direction (Fig. 3F, Fig. S7B), and polarization orientation (Fig. 3G, Fig. S7C) across the leading edge showed striking local differences. The flow direction across the two observed parts of the leading edge, lamellipodium (marked *lam* in Figs. 3E and 3F) and actin arcs (marked *arcs* in Figs. 3E and 3F) was mostly retrograde. However, the flow speed of 2-8nm/s within actin arcs (Fig. 3E, *arcs*) was substantially lower compared to the flow speed of 10-15nm/s within the actin meshwork in the lamellipodium (Fig. 3E, *lam*). The flow speeds and directions are consistent with earlier studies(7).

We observed highly oriented F-actin (in Fig. 3G, red-colored band marked *arcs*) that form the actin arcs (9) parallel to the leading edge. In contrast, the orientations of F-actin in the lamellipodium (in Fig. 3G, green-cyan-blue band marked *lam*) showed a more diverse angular distribution consistent with the meshwork architecture of actin filaments that has been studied with electron microscopy (28, 29). Our approach, for the first time, enabled direct measurement of local filament orientations relative to the flow direction (Fig. 3H) in regions of interest, such as the white dashed boxes within the lamellipodium in Figs. 3F and 3G. We observed that the flow direction within the lamellipodium is narrowly distributed (Fig. 3H, green), whereas F-actin orientation is broadly distributed, but is not isotropic (Fig. 3H, magenta). The region highlighted in Figs. 3F and 3G spans both Arp2/3-based dendritic network found within the first 1-2 μm of the leading edge and more diversely oriented filaments further away from the cell edge (28, 29). Thus, the F-actin orientation distribution we observe is consistent with data reported with electron microscopy. We did not detect a significant number of tracks that moved across the transition zone between lamellipodium and actin arcs due to the disappearance of the particles during the tracking (Video S4, Fig. S7D). However, particles observed in the transition zone have a lower polarization factor (Fig. S7F), suggesting either increased angular fluctuations in AF488-phalloidin or a broader orientation distribution of actin filaments within each particle, or both. The angular fluctuations may arise from depolymerization of actin filaments, whereas the broader distribution can arise from cross-linking actin filaments by myosin motors (9), which eventually cause contraction into actin arcs.

The above results illustrate the ability of our experimental and analysis methods to probe local order and polymerization of cytoskeletal filaments that give rise to complex micron-scale networks. Future analysis of motion, polymerization, orientation, intensity, and polarization factor as a function of distance from the leading or retracting edge promises to shed light on the organization of molecular components of the contractile actomyosin network responsible for force generation during cell migration.

### Tracking position and orientation of septin-conGFP particles in septin assemblies in living cells

We next applied our method to the septin cytoskeleton. Septins are GTP-binding proteins that self-assemble into heteromeric rods that anneal to form filaments in vitro and higher-order structures such as fibers, bundles, and rings at the plasma membrane of cells. Septin functions are diverse and range from acting as scaffolds that concentrate signaling proteins to allowing cells to perceive membrane curvature (32, 33). Atomic structural models show that mammalian septins form non-polar filaments (34). The non-polar nature of septin complexes is evolutionarily conserved and the minimal octameric rod for yeast consists of four different subunits aligned in a palindromic order Cdc11-Cdc12-Cdc3-Cdc10-Cdc10-Cdc3-Cdc12-Cdc11(35). In low salt conditions, these rods can spontaneously form long filaments and bundles. The organization of septin filaments that contained a constrained septin-GFP fusions has been studied previously by polarized fluorescence microscopy in a filamentous fungus (*Ashbya gossypii*) and in budding yeast (*S. cerevisiae*), revealing the striking rearrangement of septin filaments during division and septation (4, 5, 12, 13). The orientation of polarized fluorescence averaged over many fluorophores was found 1) to be polarized either parallel or perpendicular to the symmetry axis of the assemblies that included hourglass, ring, bar and fiber structures, and 2) to change by 90_°_ during cytokinesis. Based on our fluorescent single molecule studies of septin with non-constrained GFP expressed in *S. cerevisiae* and *S. pombe*, the minimum units in the cytosolic fraction of septins in fungi are octameric rods (36). Further, our work has shown that octameric rods and larger filaments diffuse laterally on the plasma membrane and anneal to polymerize into higher-order structures (36). However, the specific arrangement and dynamics of septin rods and filaments within the higher order structures has not been analyzed in live cells although it has been observed with localization microscopy (37).

To gain further insight into the molecular assembly of septin subunits into higher-order structures, we analyzed the polarized fluorescence of sparsely labeled septin assemblies using the instantaneous FluoPolScope. By acquiring and analyzing time-resolved polarized fluorescence, we identified the dynamics of septin-GFP particles bound to septin bars at the cell membrane of the filamentous fungus *Ashbya gossypii*. We studied the dynamic orientation of septin subunits in living cells using two different fusions of septin with constrained GFP tags (SI Notes 10 and 11) (4, 5): Cdc12-conGFP4 and Cdc12-conGFP3. The fluorescence of Cdc12-conGFP4, averaged over several fusion proteins bound to the same septin bundle, is polarized normal to the length of septin bundles (hereafter this construct is referred to as Cdc12-conGFP∥), whereas Cdc12-conGFP3, which is missing one amino acid in the protein-GFP linker compared to Cdc12-conGFP4, the ensemble fluorescence is polarized parallel to the length of the bundles (hereafter, the construct is referred to as Cdc12-conGFP∥).

Fluorescently labeled septins in *Ashbya* expressing Cdc12-conGFP∥ appeared as bright particles on bars (bundles of septins) at the hyphal cortex (5) where septa would form in later stages of cell growth (Fig. 4A). The ensemble orientation of polarized fluorescence of these particles were mostly perpendicular to the longitudinal axis of bars (Fig. 4B), as was observed in our former studies. However, septin bundles exhibit bends. Therefore, similar to the *in vitro* F-actin (Fig. 2C), we defined the local bundle orientation (*ϕ_f_*) as perpendicular to the ensemble orientation (*ϕ_e_*) averaged over the first five time-points of tracked particles.

**Fig. 4.**
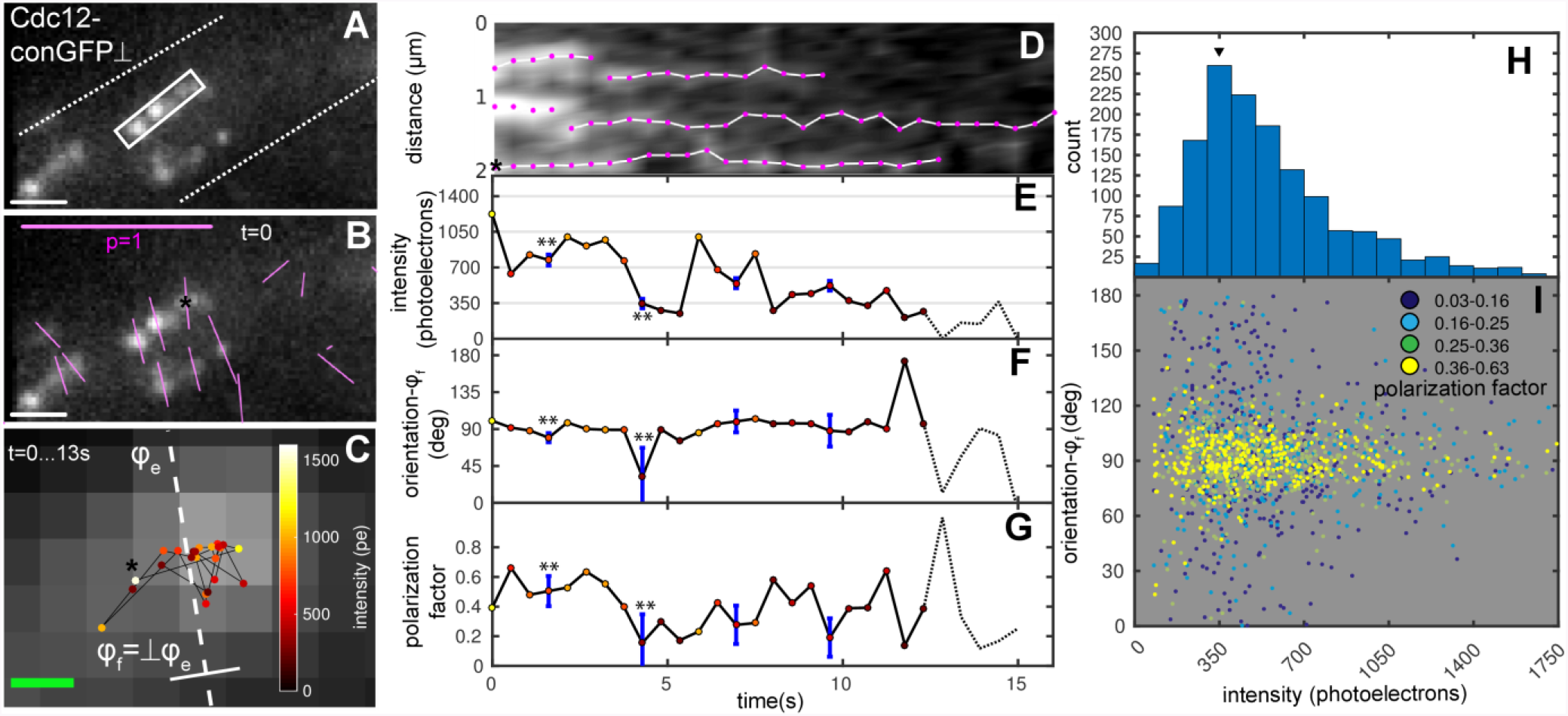
Position and orientation of single septin-conGFP⊥particles in live *Ashbya gossypii*: Data are presented in the same way as in Fig. 2. (**A**) Fluorescence images of septin-conGFP expressed in *Ashbya gossypii*. (B) The same fluorescence image in (A) with fluorescence polarization orientations indicated by magenta lines measured at the start of an image sequence. The ensemble orientation (*ϕ_e_*) of polarized fluorescence of particles is roughly perpendicular to the visible long-axis of slightly bent septin bars. The local orientation of septin bars (*ϕ_f_*) is defined to be perpendicular to local ensemble orientation. (**C**) Positions of centroid of representative particle (* in A) over time. (**D**) Kymograph of fluorescence particles along the long axis of the white rectangle in A. **(E,F,G)** Time dependent change of intensity, orientation, and polarization factor of representative particle are plotted over time in E, F and G, respectively. Blue error bars represent standard deviation due to shot-noise. This particle exhibits change in polarization factor and orientation (**) synchronous with drop to single-molecule intensity that cannot be explained by uncertainty due to shot-noise. (**H**) Histogram of particle intensities before they bleached to the background from 1200 particle observations (52 tracks). The histogram suggests that the single septin-conGFP⊥s intensity is ~350 photoelectrons. (**I**) Scatter plot of orientation (relative to *ϕ_f_*) vs. intensity (above background) of the same particles. Scale bars: white (B) is 1μm, magenta (B) is polarization factor=1, green (C) is 100nm.

We tracked the intensity, position and orientation of fluorescent particles of Cdc12-conGFP⊥ until the fluorescence signals bleached down to the background level. Fluorescent particles, whose positions were tracked for more than 10 seconds, followed a random walk within a short distance of ~350 nm (Figs. 4C, 4D, Video S6). While exhibiting highly constrained diffusive movements on the bars, the fluorescence of the particles bleached down to the background level (Fig. 4E). The histogram of fluorescence intensities of all particles (Fig. 4H) revealed that the typical intensity of a single particle was ~300-400 photoelectrons, which corresponds to the intensity level before the last bleaching step (Fig. 4E). The orientation of polarized fluorescence of Cdc12-conGFP⊥ particles was perpendicular to the bar axis irrespective of particle intensity, but occasionally deviated from this alignment by amounts that cannot simply be accounted for by the uncertainty due to shot-noise. Some of these fluctuations in orientation (^**^ in Fig. 4F) are synchronous with reduction in polarization factor (^**^ in Fig. 4G) and might be due to wobble of the fluorophores arising from displacement of the septin subunits or the flexibility of the underlying septin structure. As seen from Fig. 4I, the orientation of bright particles with a high number of septin-conGFP molecules fluctuated little around the filament orientations, while fluctuations in orientation increased at the single-molecule level. Furthermore, particles whose orientation was different from the filament orientation exhibited reduction in polarization factor.

Cdc12-conGFP∥ constructs expressed in *Ashbya* were imaged under the same conditions as Cdc12-conGFP⊥ constructs. The ensemble polarization orientation of this construct was parallel (Figs S8A,B) to the longitudinal axis of the bars as observed previously (4, 5). The intensity histogram showed a peak at 350 photoelectrons/frame (Fig. S8H). The intensity of the representative particle (Fig. S8E) fluctuated around 350 photoelectrons before bleaching to the background level. Thus, the single GFP intensity for Cdc12-conGFP∥ was consistent with the intensity for Cdc12-conGFP⊥ construct. Similar to our observations in Cdc12-conGFP⊥, the particles showed highly constrained diffusive motion on the septin bars as shown in Figs. S8C, S8D, and Video S7. The orientation of polarized fluorescence in the representative Cdc12-conGFP∥ particle shown in Figs. S8F and S8I was mostly aligned with the bar orientation, even at single molecule intensity. The polarization factor of this representative particle remained constant (Fig. S8G) at the single molecule level, in contrast to the representative Cdc12-conGFP⊥ particle shown in Fig. 4G.

We also evaluated the possibility that fluctuations of individual septin-GFP particles are caused by the movement of the underlying septin structure, the whole cell, or even drift of the microscope stage. However, since the motion of multiple septin-GFP particles in proximity of one another was observed to be asynchronous (Videos S6 and S7), we concluded that the observed motion of septin-GFP reflected the motion of individual septin filaments.

### Statistical analysis of the motion and orientation of single septin-GFP molecules on septin assemblies

We next sought to examine the constrained diffusion, both in position and in orientation, noticed for septin complexes within assemblies. To do so, we computed the mean and standard deviation of fluctuations in intensity, position, and orientation of several septin-GFP particles.

To test whether the observed fluctuations depend on the duration over which septin subunits are incorporated into the structure, we discriminated between particles (SI Note 8) already assembled into septin bars (pre-assembled particles) and those that attach to septin bars during the observation time (nascent particles). Nascent Cdc12-conGFP⊥ particles typically contained only a single GFP molecule. We analyzed pre-assembled Cdc12-conGFP⊥ particles only if their fluorescence was tracked down to the single GFP level, thereby analyzing comparable intensities between the two particle types. As a control for mobile particles with GFP, we analyzed fluctuations of a fusion protein consisting of Pleckstrin Homology (PH) domain from PLC-γ, Glutothione S-transferase (GST), and GFP expressed in *Ashbya*. This fusion protein, called PH-GFP, binds to phosphatidylinositol lipids and undergoes lateral diffusion on the cytoplasmic side of the cell membrane. Since the GST tag associated with PH-GFP dimerizes with a dissociation constant as low as 1nM (38), dimerized PH-GFP is expected to be the dominant form in our expression condition. As an immobile control, we used individual GFP particles bound to poly-D-lysine coated coverslips in our *in vitro* experiment shown in Fig. 1. Since filament orientation (*ϕ_f_*) is not applicable to PH-GFP and immobile GFP measurements, the time-resolved fluctuations are analyzed relative to the ensemble orientation (*ϕ_e_*) of each particle over first 3 observations for all three constructs.

Representative trajectories of nascent and pre-assembled Cdc12-conGFP⊥, PH-GFP bound to the cell membrane, and *in vitro* GFP bound to a poly-D-lysine coated coverslip (Fig. 5A) showed the following: PH-GFP exhibited unconstrained mobility because of the fluidic nature of the membrane. Compared to the mobility of PH-GFP, diffusive motions of both nascent and pre-assembled Cdc12-conGFP⊥ were constrained within a distance <350nm. Further, GFP bound to a coverslip were constrained within 25 nm during 50s of measurement time. Therefore, observed positional fluctuations of Cdc12-conGFP⊥and PH-GFP particles cannot be attributed to shot noise or stage drift of our imaging system.

**Fig. 5:**
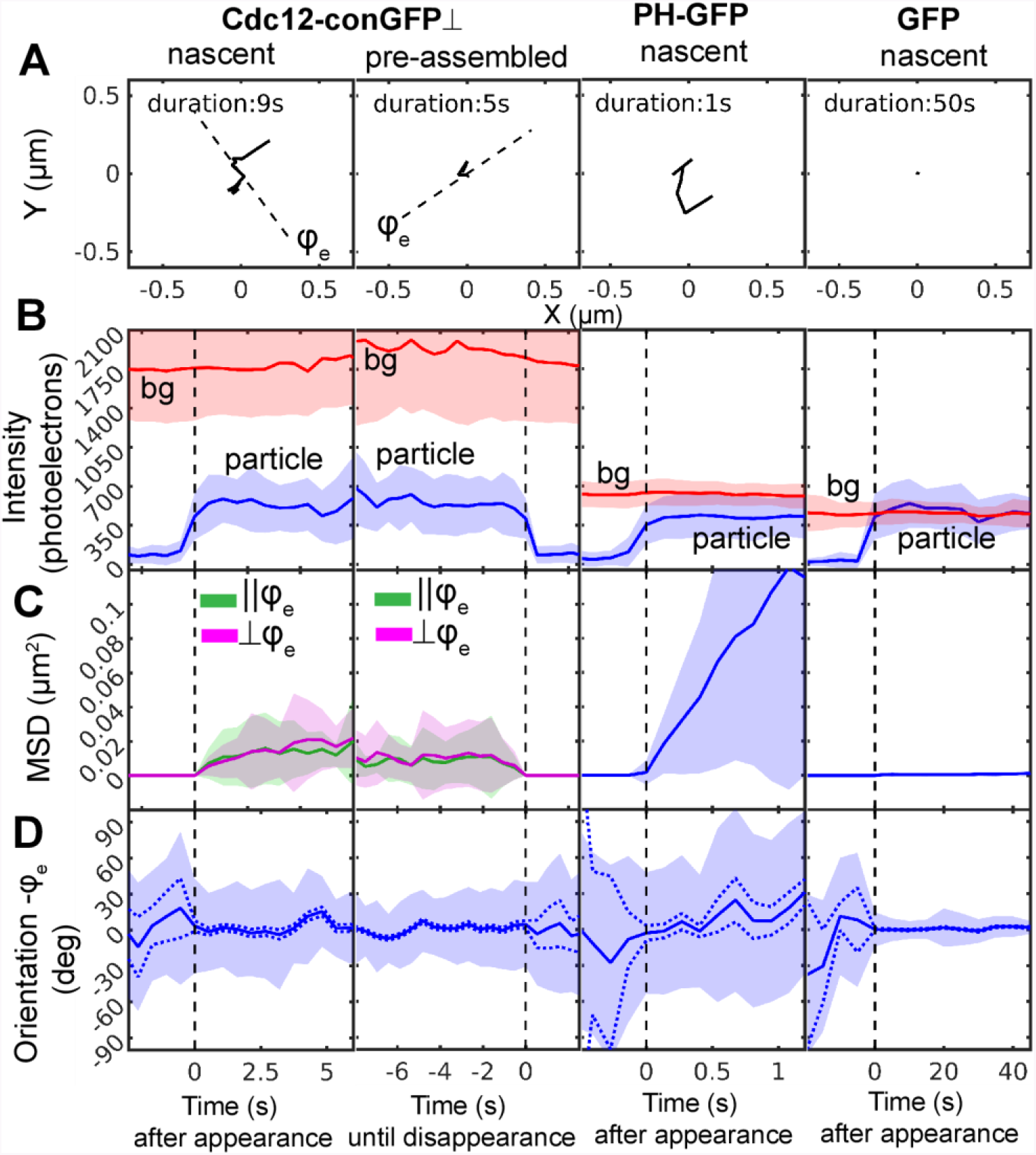
Fluctuation analysis of orientation, position, and intensity of Cdc12-conGFP⊥, PH-GFP, and GFP particles: Cdc12-conGFP⊥ were imaged continuously with 500ms exposure, 1.1 μW/μm^2^ laser power; PH-GFP were imaged continuously with 100ms exposure, 5.5 μW/μm^2^ laser power; GFP on poly-D-lysine coated coverslip was imaged at 5s intervals with 500ms exposure, 1.1 μW/μm^2^ laser power only during exposure. **(A)** Trajectories of representative particles. From left to right: nascent or newly-bound Cdc12-conGFP⊥ to septin bars, Cdc12-conGFP⊥ that were pre-assembled in septin bars, nascent PH-GFP in *Ashbya*, and nascent *in vitro* GFP bound to the surface of glass coverslips. For nascent Cdc12-conGFP, PH-GFP and *in vitro* GFP, trajectories are shown from the time they arrive within the TIR depth until their disappearance. For pre-assembled Cdc12-conGFP⊥, the trajectory is shown back in time from the time of disappearance. The dashed line in Cdc12-conGFP plots shows the ensemble polarization orientation *ϕ_e_* at the start of the acquisition. *ϕ_e_* is computed by averaging particle intensities in the first 3 frames of the track. **(B-D)**Fluctuations in intensity, position, and orientation. Fluctuations were analyzed across the ensemble of N particle tracks aligned at the frame of appearance or at the frame of disappearance. N=48 for nascent Cdc12-conGFP⊥, N=52 for pre-assembled Cdc12-conGFP⊥, N=58 for PH-GFP, and N=95 for GFP. Unlike Fig. 2I and 4I, the fluctuations are analyzed relative to the ensemble orientation (*ϕ_e_*), because filament orientation (*ϕ_f_*) is not applicable to PH-GFP and immobile GFP measurements. **(B)** Mean (solid line) and standard deviation (shaded area around mean) of particle intensity (blue) and background intensity (red) among the aligned tracks. **(C)** Mean and standard deviation of squared displacement relative the position at *t*=0. For Cdc12-conGFP⊥, the vector squared displacement is partitioned into components parallel (∥*ϕ_e_*) and perpendicular (⊥*ϕ_e_*) to the ensemble orientation. **(D)**Mean and fluctuations of polarization orientation across the aligned tracks computed using circular statistics as described in Methods. Orientation within each track is relative to its ensemble orientation as described in the legend to panel A. The shot-noise induced uncertainty in mean orientation is shown by the dotted blue line.

We next compared the intensities of nascent and pre-assembled Cdc12-conGFP⊥ particles with intensities of PH-GFP and GFP particles (Fig. 5B). In all measurements the total excitation energy per frame was the same and we found the averaged intensities were similar, within a range between 350-700 photoelectrons. PH-GFP particles exhibited fluorescence corresponding to 1xGFP (350 photoelectrons) or 2xGFP (700 photoelectrons) whose ensemble average of ~500 photoelectrons, reported in Fig. 5B (third column from the left), is consistent with the estimated number of GFP in PH-GFP particles expressed in *S. pombe* as reported in our earlier work (36). The fluorescence intensity at the location of particle appearance (Fig. 5B), measured just before the time of appearance *t=0*, was the same as the background for nascent Cdc12-conGFP⊥ and PH-GFP particles. At *t=0*, the intensity increased to one or two times the single GFP intensity. Since there are two copies of Cdc12 in a single octameric rod and the corresponding fluorescent particle should have either one or two Cdc12-conGFP, the intensity of nascent Cdc12 particles is consistent with an octameric rod that binds to the cortex from the cytosol, as previously observed *in vitro* (36).

To analyze fluctuations in position, we computed squared displacements of particles relative to their position at the time of appearance for nascent particles, or relative to their position at disappearance for pre-assembled particles (Fig. 5C). The squared displacement of Cdc12-conGFP ⊥ particles was resolved along directions parallel (∥*ϕ_e_*) and perpendicular (⊥*ϕ_e_*) to the ensemble polarization orientation of each particle. The mean squared displacement (MSD) of PH-GFP particles increased over time with a nearly constant slope, suggesting simple 2D diffusion over the time period of 1s. The MSD of GFP particles stays near zero for 60s, confirming their immobile attachment to poly-D-lysine coated glass coverslips and the stability of the microscope setup. The MSD of both nascent and pre-assembled Cdc12-conGFP⊥ particles exhibited a slope that reduces over time, suggesting confined diffusion. Resolving the motion of the Cdc12-conGFP⊥ particles parallel and perpendicular to the length of septin assembly did not provide evidence for a dominant direction of diffusive motion with respect to the orientation of the assembly over 6s duration.

To validate our finding of positional fluctuations of septin-GFP particles, we analyzed positional fluctuations of another septin-GFP construct, Cdc11-GFP, expressed in *Ashbya.* In this construct, GFP is not constrained in orientation with respect to Cdc11, another subunits in the septin octamer (33). We found that fluctuations in intensity and position of Cdc11-GFP particles were nearly identical to Cdc12-conGFP, further confirming that the minimum unit of assembly is an octamer (Fig. S9B) and that septin octamers undergo constrained diffusion in the larger assembly indicating this behavior is not a consequence of constraining the fluorescent tag (Fig. S9C).

We estimated the size of a ^‘^corral^’^ that confines septin-GFP and the diffusion coefficient within the corral, using a widely-used model of confined diffusion of membrane receptors (39). Using this model, we estimated the diffusion coefficient *D (μm^2^s^-1^)* within the corral by fitting a line to MSD vs. time data (*t*) near *t=0* (Fig. 5C, Fig. S10) according to the relationship *MSD=4Dt*. The corral size (*L*) and the asymptotic value of MSD (*MSDmax*) are related by *L2/6*=*MSD_max_*. As shown in Fig. S10, *D=*0.03 *μm^2^s^-1^* for PH-GFP, whereas *D=*0.003 *μm^2^s^-1^* for pre-assembled Cdc12-conGFP and Cdc11-GFP particles. Note that we added MSD along and perpendicular to the ensemble orientation together to estimate the diffusion coefficient of Cdc12-conGFP. Thus, our data indicate that septin-GFP particles experience confined diffusion that is an order of magnitude slower than diffusion experienced by PH-GFP. The corral size (*L*) for Cdc12-conGFP⊥ was estimated to be 350nm and for Cdc11-GFP estimated to be 250nm. With the caveat that the chosen motion model is the first approximation of the motion of septin-GFP, we infer that the lateral extent over which the septin-GFP can move near the binding site is an order of magnitude longer than the length of a single septin octamer (32nm).

We calculated the mean orientation and the spread of the orientations (SI Note 12, Fig. S11) across selected tracks as a function of time, as well as shot-noise induced uncertainty in mean orientation (SI Note 5). The polarization orientation of pre-assembled and nascent Cdc12-conGFP⊥ particles (Fig. 5D) remained along the ensemble fluorescence polarization of the underlying septin bars. However, the orientations fluctuated in time, exhibiting a large spread (~30^°^, blue patch in Fig. 5D left) that cannot be explained by uncertainty due to shot-noise (dashed line in Fig. 5D left). This finding indicates that the orientation of Cdc12-conGFP⊥ particles itself fluctuates in time while bound to the septin bars. On the other hand, the orientation of Cdc11-GFP before and after binding fluctuates widely (Fig. S9D) since GFP was not constrained to Cdc11. The orientation of PH-GFP particles also varied widely and once bound to the membrane fluctuated randomly. For GFP molecules that were bound to the surface of poly-D-lysine coated glass coverslips, angular fluctuations after binding were less than 10^°^.

To check if the fluctuations in orientation and position of septin-conGFP are consistent after repeated imaging of the same structure, we bleached the same septin bars up to 5 times, each time followed by a recovery period of 2-10 minutes (Fig. S12). The structural integrity of septin bars after repeated bleaching and observation of fluctuations in the orientation between two septin-conGFP constructs indicate that the observed dynamics of GFP dipoles reflect bona fide dynamics of septin subunits and not artifacts induced by imaging.

The above observations identify two remarkable features of septin-GFP molecules bound to micron-scale septin assemblies, when analyzed with the instantaneous FluoPolScope. First, the Cdc12-conGFP and Cdc11-GFP particles exhibited confined movement in position that is an order of magnitude slower than diffusion of the model peripheral membrane protein PH-GFP, while the linear range of motion is an order of magnitude larger than the length of a single septin octamer. Second, the ensemble mean orientation across many septin-conGFP particles remained consistent as they bleached from multiple to single fluorophores, albeit exhibiting fluctuations in the orientation larger than possible only from shot-noise. The consistency of polarized fluorescence orientation even down to a single fluorophore is compatible with, but not predicted by the earlier studies of ensemble polarized fluorescence from densely labeled septin structures (5, 13). Fluctuations in both position and orientation of septin-GFP can arise from either the flexibility of polymerized septin filaments or from the movement of the subunits within the polymeric scaffold. We speculate that the movements we see may be due to a combination of fragmentation and annealing events of filaments within and between filaments that are bundled together into a septin bar. Further analysis of motion, intensity, orientation, and polarization factor of septin-GFP constructs in live cells over longer time-scales has the potential to address the question of how septin assemblies form the observed diversity of structures and how they can undergo large-scale rearrangements during cytokinesis (4, 5).

## Conclusion

We are reporting the design, implementation, and application of an instantaneous Fluorescence PolScope, employing isotropic total internally reflected (TIR) excitation and instantaneous imaging along four equally spaced polarization orientations. The key advantage of our method over earlier methods is its ability to report molecular orientation of dynamic assemblies of arbitrary symmetry and its robustness to background fluorescence typically observed in live cells. High background fluorescence can limit the accuracy of molecular orientation measurement in living cells as clarified in the paper. The robustness to background fluorescence is achieved using total internal reflection excitation and single particle analysis algorithms that take into account remaining spatially variable background. The instantaneous acquisition and temporal tracking of intensity, position, and orientation of fluorescently labeled molecules allows us to study structural dynamics of mobile molecules at 10Hz. This computational analysis should enable study of variety of questions that require dissection of sub-resolution molecular order, including structural organization of DNA in chromosomes, morphological changes in cellular membrane, and dynamic assembly/disassembly of the cytoskeletal networks. Our method is still limited by its inability to report the tilt angle of dipoles (Z-axis component of the dipole distribution), because polarization-resolved measurement within the XY plane is sensitive only to the projection of polarized fluorescence on the focal plane. This limitation will be overcome in the future by rapid switching of the intensity of the Z-axis component of polarized excitation while acquiring the polarization-resolved emission.

We used the instrument and the complementary algorithms to analyze the time-resolved orientation of single molecules *in vitro* and in living cells. *In vitro*, we demonstrated the orientation of individual TOTO-1 fluorophores that intercalated into combed DNA and whose fluorescence is polarized perpendicular to the DNA axis, consistent with the known atomic structure of the complex. Analysis of *in vitro* actin filaments labeled with AF488-phalloidin showed that the polarization orientation of even a single fluorophore reliably reports the local orientation of the actin filament to which it is bound. By sparsely labeling actin network with AF488-phalloidin, we provided a first demonstration of analysis of local orientation of actin filaments relative to retrograde flow at the leading edge in live cells. Our findings are consistent with previous studies of actin filament orientation using electron microscopy and actin network flow using fluorescent speckle microscopy. The 100 millisecond temporal resolution, 25nm localization accuracy, and shot-noise limited accuracy of 3^°^ in the measured orientation of single molecules allowed us to analyze dynamic position and orientation of individual septin proteins during their interactions with larger assemblies in living cells. Our single molecule position analysis revealed that septin octamers undergo a confined random walk, which is an order of magnitude slower than other membrane-associated proteins. Moreover, the orientation analysis revealed that the orientation of septin-GFP fluctuates around the net orientation of the underlying septin assembly.

## Methods

### Isotropic total internal reflection excitation

We employ a circularly polarized laser beam that is rotated in the back focal plane of the objective (18) with rotation frequency higher than 300Hz to achieve isotropic illumination at the specimen plane (Fig. 1B, methods). We used total internal reflection (TIR) to produce isotropic 3D excitation as illustrated in Fig. 1B. The radius of the annulus in the BFP was adjusted such that the collimated laser reached the specimen plane beyond the critical angle required for TIR. Circularly symmetric TIR excitation with circularly polarized light leads to equal amounts of p- and s-polarized excitation over 360o. Integrating the evanescent field components along the X, Y, and Z axes under this symmetric illumination scheme shows that the intensity of the field component along the Z-axis is ~8 % less than the intensity components along X and Y. A laser beam (DPSS laser BCD, 488nm, Melles Griot) was circularly polarized by a quarter wave plate (WPQ-4880-4M, Sigma-koki) and focused into the back focal plane (BFP) of the objective lens (PlanApo TIRF 100x 1.49NA oil, Nikon) on an inverted microscope (TE2000-E, Nikon). The laser intensity at the specimen plane was 5.7μW/μm^2^ when no ND filter was inserted into the light path. The beam was scanned along a circular annulus in the BFP using a pair of galvo-scanned mirrors (GVS202, Thorlabs).

### Instantaneous polarization analysis along four orientations

Polarization analysis in the aperture space ensures that fluorophores are subjected to identical optical analysis irrespective of their position in the specimen plane. We do so (Fig. 1B) by re-collimating the light originating from the primary image plane of the microscope, splitting the emission according to polarization, and finally focusing onto the four quadrants of an EMCCD camera (iXon+, Andor Technology). In the collimated space, the fluorescence is first divided equally between two arms by a polarization independent beam splitter (50/50 in Fig. 1B, 21014, Chroma Technology). A pair of polarization beam splitters (PBS in Fig. 1B, Semrock) separates the fluorescence into four images and analyzes their linear polarization along 0^°^, 45^°^, 90^°^ and 135^°^ orientations. One PBS generates 0^°^ and 90^°^ polarization beam paths, while the polarization in the other arm is first rotated by 45^°^ using a half-wave plate (HWP in Fig. 1B, Meadowlark Optics NH-532) and then passed through the other PBS to generate the 45^°^ and 135^°^ paths. Subsequently, we use broadband mirror-based beam-steering optics and a focusing lens (FL in Fig. 1B) to project four images onto the four quadrants of a single EMCCD detector. The collimating and focusing lenses together magnify the image by 1.5x. We acquire data using a 100x 1.49NA objective lens and 1.5x tube-lens, leading to a total magnification of 225x between the specimen and the camera. The width of the camera pixel is 16 μm leading to the pixel-size at the specimen plane of 70nm, which provides slightly better than Nyquist sampling of the point spread function.

### Registration of four quadrants and intensity calibration

All algorithms were developed in MATLAB and are available upon request.

To register the quadrants, we recorded images of fluorescent latex beads (diameter, 0.12 μm, Life technologies) spread on the surface of a glass coverslip (Fig. S2A). After cropping raw camera frames into quadrants, intensity-based optimization (MATLAB’s imregister function) was used to compute affine transformations that minimize intensity differences among quadrants. A sensitive readout of achieving sub-pixel registration between the four quadrants is the disappearance of a systematic polarization pattern for a sample like the latex beads that must not show any polarization. As shown in Fig. S2B, our approach achieves registration with sub-pixel accuracy.

To normalize differences in transmission efficiency between the four imaging paths, we imaged a 20 μM fluorescein solution excited by uniform, isotropic polarization illumination from a 100W tungsten halogen lamp through an excitation filter (Semrock FF02 473/30). Using the isotropic nature of fluorescence polarization from a solution of randomly oriented fluorescein molecules, we generated correction images that when multiplied with raw sample images, compensated the position-dependent differences of the transmission efficiency in the four-quadrant polarization optics.

Subsequently, we calibrated polarization-dependent intensity differences due to unbalanced extinction factors of the polarization optics using the approach of calibrating an industrial polarimeter (40). In this approach, polarized light at several known states is presented to the optical system and recorded intensities are analyzed to compute an ‘instrument matrix’ that converts Stokes parameters of known polarization states to recorded intensity. As a specimen with known pattern of polarization transmission, we used a tangential polarizer (Photonic Lattice Inc., Japan), whose polarization transmission axis is tangentially aligned along coaxial circles (Fig. S2C, SI Note 1).

By using these three calibration objects, namely fluorescent beads, a fluorescein solution and a tangential polarizer, we have obtained registration with sub-pixel accuracy, correction factors for compensating transmission differences, and balanced the extinction differences of the four polarization-sensitive light paths.

Finally, we calibrated isotropic TIR excitation by minimizing the fluorescence polarization from immobilized fluorescent beads absorbed to the surface of glass coverslips, as any biased polarization in the illumination caused a fluorescence polarization signal from immobilized fluorescent beads. We manually tuned the half-wave plate and quarter-wave plate that were located in the excitation illumination path so that the polarization factors of imaged fluorescent beads were minimized.

### Computation of orientation, polarization factor, and intensity

The polarization of a fluorescent particle was characterized by its degree of polarization, called the polarization factor (*p*), and the orientation (*ϕ*) of its maximum polarization. The particle intensity recorded in each quadrant is related to the average intensity across the four quadrants (*I*), polarization factor, orientation, and background intensity (*Ibg*) as follows (22):

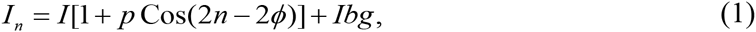

where,*n* = 0^°^, 45^°^, 90^°^, 135^°^.

Retrieval of the orientation and polarization factor of particles is efficiently expressed in terms of the Stokes parameters of the polarization-resolved emission:

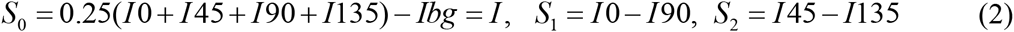

While the estimates of *S*_1_, *S*_2_ are independent of the isotropic background intensity *Ibg*, their uncertainty is impacted by the shot-noise of the background.

The above relationship can be written in matrix form as **S** =*M*(**I** -*Ibg*), where, **I**=[*I*0, *I*45,*I*90,*I*135]*T* is the column vector of input intensities, **S** =[*S*0, *S*1, *S*2]^T^ is the column vector of Stokes parameters, and *M*, called instrument matrix, is the matrix of coefficients that relate these two vectors according to Eq. 3. The matrix *M* in above equation is replaced by the instrument matrix that is generated using the tangential polarizer as described in SI Note 2.

The total particle intensity (in photoelectrons), polarization factor (dimensionless between 0 and 1), and ensemble orientation (in radian) are obtained from Stokes parameters as follows:

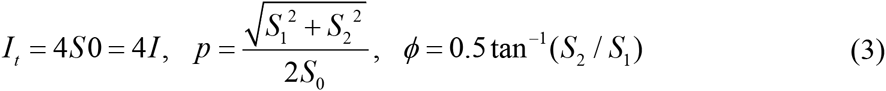

During a camera exposure, a range of dipole orientations is observed due to several factors: rotational wobble of a single dipole due to flexible linkers to the structure of interest, or motion of the structure itself in live cells.

## Acknowledgements

The authors acknowledge helpful discussions with Andrew Bridges, Hari Shroff, Vinay Swaminathan, Clare Waterman, Michael Shribak, Takayuki Nishizaka, Takeharu Nagai, and many of our summer collaborators at the MBL. This work was supported by grants from the National Institutes of Health to TT (R01 GM100160), to RO (R01 GM114274), and to MM (T32 GM008704); from the National Science Foundation to ASG (MCB 1212400); from the Human Frontier Science Program to SBM (LT000096/2011-C); from the Marine Biological Laboratory to TT (Start-up funds from Inoue Family Endowment), to SBM (2014 Neal Cornell Career Development Award, 2015 MBL-UChicago collaboration award), and to ASG (Whitman summer investigator award). The authors are grateful for Japan Science and Technology Agency PRESTO program, which had supported the early phase of this research and generously offered us essential optical instruments.

